# The Syncytial Mesh Model: A Mesoscale Control-Field Framework for Scale-Dependent Coherence in the Brain

**DOI:** 10.1101/2024.11.22.624908

**Authors:** Andreu Ballús Santacana

## Abstract

Large-scale neural coherence and distributed plasticity remain only partially accounted for by circuit- and connectome-based models: stable phase gradients, scale-dependent harmonic structure, and non-local functional reorganization frequently span regions lacking direct synaptic linkage. We propose the Syncytial Mesh Model (SMM) as a *candidate* mesoscale framework in which astrocytic syncytia operate as a slow *control-field substrate* that *shapes*—rather than directly generates— the dynamical geometry within which neuronal populations evolve. The SMM is articulated as a three-layered system: (i) local neural circuits, (ii) structural connectomic pathways, and (iii) a continuous mesoscale field grounded in astrocytic network physiology. The third layer is implemented as a damped wave equation on a small-world astrocytic topology and is presented as a *phenomenological effective theory*, not as a microscopic biophysical model. Numerical simulations— using a 9-point isotropic Laplacian, perfectly matched layer (PML) boundaries, and unified RK4 integration—produce illustrative phenomenological dynamics: artifact-free amplitude snapshots, radial phase gradients, and low-frequency mode selection that are consistent with, but not exclusive to, delta/theta-band features reported in human MEG and LFP. An analytic two-mode coherence model fitted to phase-gradient coherence across *N* = 43 subjects yields a scaling parameter *λ*_0_ ≈ 1.5903*/*s producing a plateau at ~ 4.65 % for millimeter-scale patches. The comparison of simulated and empirical spectra (median Pearson *r* = 0.917, median MSE = 26.6 dB^2^) is offered as *proof-of-principle phenomenological compatibility*, not as empirical confirmation of the model. We position SMM relative to neural-field, connectome-harmonic, metastability, and predictive-processing frameworks, and we propose four falsifiable discriminatory predictions that distinguish SMM from alternatives. The strongest claim of this framework is not that astrocytes generate cortical rhythms, but that astrocytic syncytia plausibly provide a slow dynamical control geometry for large-scale neuronal organization.

## 1 Introduction

The emergence of large-scale synchrony and distributed plasticity is central to cognition, yet several experimentally documented features are difficult to account for using neural-mass and connectome models alone. First, phase gradients and traveling waves traverse non-contiguous cortical regions lacking direct synaptic connections [1, 2]. Second, low-frequency spectral features in the delta/theta range (1 Hz to 8 Hz) display considerable cross-subject and cross-species stability that is only partially explained by anatomical variability [3, 4, 5]. Third, non-invasive stimulation (TMS, tACS) and spontaneous plasticity induce functional reorganization in regions lacking direct structural links [6, 7, 8]. These observations are compatible with the existence of an additional substrate operating in parallel with canonical neural circuitry.

We use the term *phase coherence* to refer to sustained, non-zero phase offsets across space—distinct from strict zero-lag synchrony or amplitude correlation. Cortical coherence typically decays over a few centimeters, beyond which synaptic coupling alone offers a limited account [9]; yet traveling delta-band waves often span entire hemispheres, motivating the search for complementary mechanisms. (Throughout, we use “mode” for spatial eigenfunctions of the Laplacian operator, “coherence” for cross-site phase stability in field oscillations, and “resonance” for selective amplification of modal components.)

We propose the Syncytial Mesh Model (SMM) as one candidate mesoscale mechanism, in which a continuous mesh-like field—grounded in astrocytic syncytia physiology—coexists with local neural masses and anatomical connectivity. Astrocytic networks form small-world syncytia via gap junctions, propagate calcium signals at 10 µm/s to 30 µm/s, and modulate synaptic function across distances [10, 11, 12]. We do *not* claim that this layer generates the measured electromagnetic signals recorded in EEG or MEG; neurons remain the dominant current generators. Rather, the SMM proposes that the syncytial substrate *shapes the dynamical geometry* of large-scale neural organization: it constrains synchronizability, biases metastable transitions, and modulates the gain landscape in which neuronal rhythms evolve.

In this article we:

- Articulate why a slow glial mesoscale substrate is a plausible candidate—rather than a necessary one—for the phenomena above (Section 2).
- Derive the Syncytial Mesh layer as a damped wave equation on a 32 mm × 32 mm grid with a 9-point isotropic Laplacian, PML boundaries, and unified RK4 integration, framed explicitly as a phenomenological mesoscale effective theory (Sections 3–4).
- Present illustrative dynamical structures—artifact-free amplitude snapshots (Fig. 2), radial phase gradients (Fig. 3), and low-frequency mode selection (Fig. 4)—together with an analytic two-mode coherence model fitted to phase-gradient coherence across *N* = 43 subjects (Figs. 5–6) (Sections 5–6).
- Compare the SMM with thalamocortical, connectome-harmonic, neural-field, metastability, and predictive-processing frameworks, positioning the SMM as a *complement* rather than a substitute (Sections 7–8).
- Articulate four falsifiable discriminatory predictions that distinguish SMM from alternative accounts (Section 9).

The objective of this paper is to articulate a falsifiable, mechanistically grounded *candidate* mesoscale substrate. The SMM is not advanced as the explanation of cortical coherence; it is advanced as one specific, mathematically tractable, biologically anchored hypothesis that can be tested against, and combined with, alternative frameworks.

## 2 Why a Glial Mesoscale Layer?

Several candidate substrates could in principle organize slow, spatially distributed coherence beyond local synaptic coupling. Volume conduction is rapid but spatially diffuse and lacks state-dependent information storage; neuromodulator diffusion is slow but topologically weakly structured; thalamocortical loops are anatomically constrained and operate at faster timescales; connectome-harmonic accounts [48] capture structural eigenmodes but are static with respect to dynamical state. None of these alternatives is incompatible with the SMM; the question is what astrocytic syncytia could distinctively contribute.

Astrocytic syncytia combine several features that, taken together, are unusual:

- *Slow integration timescales* of seconds, well matched to infra-slow and delta/theta band dynamics [12, 17].
- *Spatial continuity*: overlapping astrocytic domains tile cortical tissue without the discreteness of axonal projections [13, 49].
- *Gap-junction-mediated electrical continuity*: connexin 43/30 coupling supports syncytial isopotentiality and quasi-continuous field organization over centimeters [16].
- *Metabolic gating*: astrocytic regulation of glucose and lactate supply directly modulates the energetic landscape of local circuits [49].
- *Neuromodulatory sensitivity*: astrocytes respond to monoamines, acetylcholine, and noradrenaline, suiting them to state-dependent gain modulation [12, 20].
- *State persistence*: Ca^2+^ dynamics carry information across hundreds of milliseconds to seconds, supporting slow control rather than rapid generation.

We do not claim that these features are unique to glia, nor that astrocytes are the only substrate capable of mesoscale control. We claim only that this constellation of properties makes astrocytic syncytia a plausible, biologically grounded candidate substrate for the kind of slow control-field dynamics articulated in the remainder of the paper.

## 3 Biological Substrate: Astrocytic Syncytia as a Mesoscale Control Field

Astrocytic syncytia form a continuous, small-world network via gap junctions (connexin 43/30), tiling cortical and subcortical territories with overlapping domains [13, 14, 52]. Two-photon imaging and connectomic reconstructions report high clustering coefficients (0.4–0.7) and short path lengths, optimizing both local integration and long-range coupling [15, 11]. White-matter astrocytes exhibit anisotropic connectivity that could support directional propagation [13]. These properties are consistent with syncytial isopotentiality, whereby extensive gap-junction coupling maintains relatively uniform field potentials over millimeter-to-centimeter scales [16].

### 3.1 Wave Propagation and Modal Structure

Astrocytic networks transmit traveling calcium signals at speeds *c* ≈ 10 µm*/*s to 30 µm*/*s, with characteristic delays of 3 s to 10 s over 100 µm domains [10]. Under appropriate boundary conditions, the resulting field admits standing modes whose modal structure is broadly in the low-frequency range compatible with delta/theta features observed in MEG/EEG [18, 5, 4]. In an idealized 1D approximation,

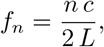

where *L* is a characteristic domain length. This expression should be read as a *heuristic modal intuition* and *phenomenological scaling estimate*, not as a biologically exact prediction of cortical resonant frequencies. Real cortex is anisotropic, heterogeneous, curved, and nonstationary, and admits a continuum of overlapping modes rather than discrete cavity resonances. 2D/3D generalizations yield similar qualitative scaling [3].

### 3.2 Astrocytes and Mesoscale State Modulation

Astrocytes modulate neuronal excitability via gliotransmitter release (glutamate, D-serine, ATP), neurotransmitter uptake, K^+^ buffering, and synaptic modulation at the tripartite synapse [12, 19, 20]. These mechanisms allow astrocytic activity to bias the timing and amplitude of low-frequency oscillations across distant populations, providing a candidate substrate for state-dependent modulation of phase coherence [21, 22, 23]. In vivo imaging shows astrocytic Ca^2+^ waves often precede large-scale neural state transitions [21, 24], and optogenetic manipulations are consistent with a causal role in slow cortical state changes [12].

This evidence supports a *control-field* interpretation: the astrocytic layer shapes the dynamical landscape (gain, damping, metastability) on which neuronal oscillations evolve, rather than itself generating the electromagnetic signals recorded at the scalp.

### 3.3 Distributed Plasticity and Synaptic Co-Activation

Astrocytic calcium signals can gate synaptic plasticity by modulating LTP/LTD induction thresholds, spatially distributing synaptic tagging signals, and coordinating cross-synaptic updates [25, 26, 27, 28, 46]. Blockade of astrocytic Ca^2+^ signalling has been reported to disrupt Hebbian learning and non-local synaptic changes [28, 12], consistent with the mesh-driven plasticity contribution in our model. Astrocytic plasticity itself—via receptor density, gap-junction permeability, or calcium response thresholds—could dynamically reshape the mode structure of the control field, providing a possible substrate for slow, state-dependent learning.

### 3.4 Empirical Parameter Bounds

Experimental measurements constrain plausible Syncytial Mesh parameters (see Table 1): wave speed *c* = 10 µm*/*s to 30 µm*/*s, propagation delay (100 µm) = 3 s to 10 s, domain size *L* = 0.1 cm to 2 cm, clustering coefficient = 0.4–0.7, Ca^2+^ wave decay time = 1 s to 10 s, gap-junction density ~10^3^/cell [10, 15]. These bounds inform simulation and analytic modeling without committing the model to specific microscopic values.

**Table 1.**
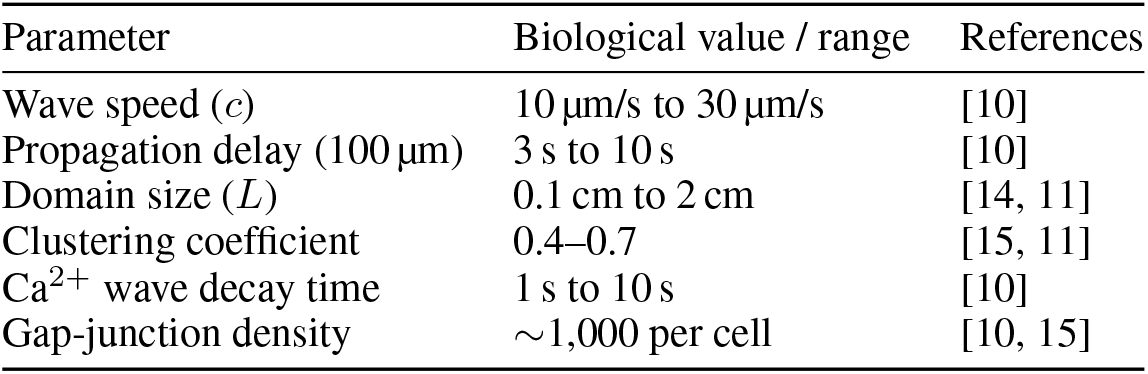
Empirically constrained parameter ranges used in the Syncytial Mesh layer.

### 3.5 Biophysical Approximations and Their Scope

Gap-junction conductance is inherently nonlinear (e.g., Cx43 exhibits voltage-dependent gating), which matters for long-distance regenerative Ca^2+^ propagation; our linear damped wave PDE approximates this behaviour only to first order [30]. Syncytial topology significantly influences wave propagation and coherence patterning, in some regimes more strongly than intracellular parameters [31, 32]. The present formulation assumes effectively uniform coupling, and future work could incorporate structured or modular gap-junction geometries, regional heterogeneity in *γ* and *c*, and explicit nonlinear reaction-diffusion terms.

### 3.6 Domain of Applicability

The Syncytial Mesh substrate, as formulated here, is primarily intended to apply to infra-slow, delta, and theta dynamics (*<* 12 Hz). Faster gamma/beta synchrony arises primarily from fast GABAergic circuits and direct axonal coupling [33, 34, 35] and lies outside the scope of this model. Astrocyte density and domain overlap vary across regions, so mesoscale effects are expected to be most relevant in associative cortex and attenuated in highly myelinated areas [13, 14]. Astrocyte syncytia develop gradually during the postnatal period [36, 53], and their alteration in aging or disease may contribute to observed shifts in slow coherence structure.

### 3.7 Summary

Astrocytic syncytia—characterized by small-world topology, slow propagation, and multimodal coupling—possess properties consistent with the kind of mesoscale control substrate articulated by the SMM. Their active, state-dependent dynamics make them a plausible, though not necessarily unique, substrate for slow, scale-dependent control of phase coherence and distributed plasticity [37, 12, 23].

## 4 Theoretical and Mathematical Framework

The SMM posits that brain-wide coherence and plasticity reflect the coupled dynamics of three interacting layers:

i. local neural circuits,
ii. macrostructural connectomic pathways, and
iii. a continuous mesoscale field plausibly implemented in astrocytic syncytia, treated here as an *effective phenomenological theory*.

Each layer is formalized below. The third-layer PDE is not a microscopic biophysical reconstruction; it is a mesoscale effective description whose virtue lies in its tractability and in admitting falsifiable predictions.

### 4.1 Layer 1: Local Neural Circuit Dynamics

Each brain region *i* is modeled with a Wilson–Cowan-type neural mass [39], in the broader tradition of coupled neural-mass models exemplified by Jansen and Rit [38]:

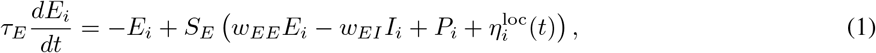

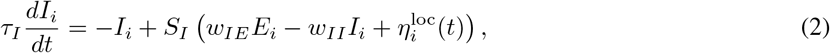

where *w*_*ab*_ are synaptic weights, *P*_*i*_ is external or inter-regional input, and 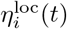 is local noise. The activation functions are sigmoidal:

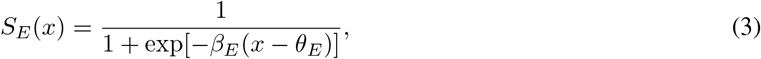

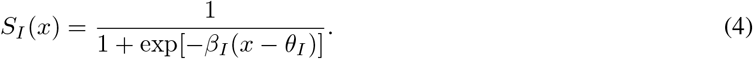

This layer generates fast oscillations and local circuit dynamics of the type observed in M/EEG and LFP recordings.

### 4.2 Layer 2: Structural Connectome and Long-Range Coupling

Anatomical pathways are represented by a symmetric connectivity matrix *C*_*ij*_, derived from diffusion MRI. Inter-regional phase dynamics follow a generalized Kuramoto model augmented by the Syncytial Mesh field:

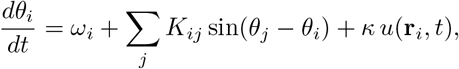

where *θ*_*i*_ is the phase of region *i, ω*_*i*_ its intrinsic frequency, and *u*(**r**_*i*_, *t*) the local mesoscale field. Coupling strengths are normalized as *K*_*ij*_ = *k*_0_*C*_*ij*_*/* Σ_*j*_ *C*_*ij*_. This layer supports network-level synchronization, hub formation, and global oscillatory modes consistent with empirical findings.

### 4.3 Layer 3: Syncytial Mesh as Phenomenological Mesoscale Field

The Syncytial Mesh substrate is described by a continuous field *u*(**r**, *t*) governed by a damped wave PDE:

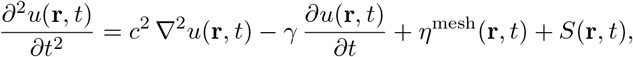

where *c* is an effective propagation speed, *γ* a damping coefficient, *η*^mesh^ space–time white noise, and *S*(**r**, *t*) a source term encoding neural-to-astrocyte coupling. Boundary conditions are implemented via a perfectly matched layer (PML) to prevent spurious reflections. The PDE is a mesoscale effective description that approximates the syncytium’s coarse-grained behaviour and connects naturally to the broader literature on neural field theory [54, 55]; it is *not* a microscopic model of astrocyte biophysics.

This field is expected to support, qualitatively:

- Smooth phase gradients and slow traveling structures less constrained by synaptic connectivity than purely network-based models.
- Low-frequency modal selection of the form *f*_*n*_ ~ *nc/*(2*L*) (1D heuristic), broadened and distorted in realistic geometries.
- Distributed co-activation patterns that bias plasticity beyond direct anatomical links.

### 4.4 Plasticity Rules: Local and Mesh-Driven Components

Synaptic plasticity is modeled as a combination of Hebbian / spike-timing-dependent updates and mesh-driven co-activation:

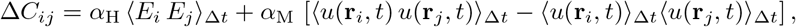

with temporal averaging over 1 s to 10 s (astrocytic integration timescale). The relative weights *α*_H_ and *α*_M_ control the balance between purely synaptic and mesoscale-modulated learning.

### 4.5 Cross-Layer Coupling

- Local neural activity (*E*_*i*_, *I*_*i*_) drives *S*(**r**, *t*) through tripartite-synapse astrocytic transduction.
- The mesoscale field *u*(**r**, *t*) *modulates* neural excitability and phase dynamics (*E*_*i*_, *θ*_*i*_); it does not replace or override them.
- Plasticity updates integrate Hebbian and mesh-driven contributions, reflecting dual synaptic and glial influences.

The field can be represented on a continuous lattice or via a graph Laplacian over empirically derived astrocytic networks. Inhomogeneities and boundary conditions can be tuned to anatomical data.

### 4.6 Summary

The SMM combines: (i) neural-mass dynamics for local excitatory/inhibitory populations; (ii) empirically grounded connectome-based Kuramoto phase coupling; (iii) a phenomenological mesoscale field on a small-world astrocytic substrate; and (iv) dual plasticity rules merging Hebbian and mesh-driven mechanisms. The system permits bifurcation analysis, efficient simulation, and parameter inference from neurophysiological recordings, offering a testable candidate framework for slow, scale-dependent coherence and distributed plasticity.

## 5 Methods

All simulations, analytic calculations, and empirical comparisons were performed in Python (NumPy, SciPy, Matplotlib, Pandas) within Google Colab. Two complementary pipelines were used. Simulation and analytic outputs were compared with empirical neurophysiological data from human subjects (MEG/EEG); these comparisons are reported as proof-of-principle phenomenological compatibility, not as empirical confirmation of the model:

- Numerical simulation of the Syncytial Mesh layer (Section 5.1): produces mesh-field amplitude snapshots, phase-gradient maps, and power spectra, compared qualitatively and statistically with empirical M/EEG features.
- Analytic two-mode coherence model (Section 5.3): parameterized and fitted to empirical distributions of phase-gradient coherence.

Supplementary materials (code, raw data, auxiliary figures) are described in Section 5.4.

### 5.1 Mesh Simulation

#### 5.1.1 Domain, Discretization, and Physical Parameters

- Spatial domain: 32 mm × 32 mm patch, discretized on a 32 × 32 grid (Δ*x* = Δ*y* = 1 mm).
- Temporal domain: *t* ∈ [0, 30 s] with Δ*t* = 1 ms (*N*_*t*_ = 30,000 steps).
- Effective wave speed: *c* = 0.015 mm*/*ms = 15 mm*/*s.
- Background damping: *γ*_bg_ = 0.10*/*s in the interior.
- PML boundaries: border of 4 grid points (4 mm) on each side; within PML, *γ*_pml_ = 2.00*/*s absorbs outgoing waves.
- Fields: *u*_*i,j*_(*t*) = mesh amplitude, *v*_*i,j*_(*t*) = *∂u*_*i,j*_*/∂t* = mesh velocity.

#### 5.1.2 Governing Equations and Discretization

The Syncytial Mesh field satisfies the effective mesoscale PDE:

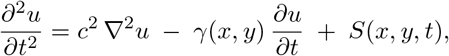

rewritten as:

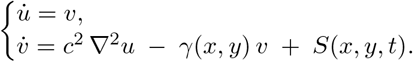

The discrete Laplacian uses a 9-point isotropic stencil:

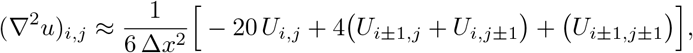

which enforces a smooth, rotationally symmetric approximation. PML handles boundary absorption without explicit periodicity.

#### 5.1.3 Stimulus Profile

For 0 ≤ *t* ≤ 1 s:

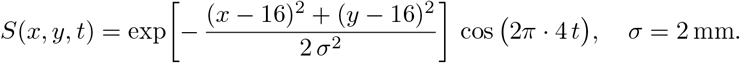

After *t >* 1 s, *S* = 0. Centered at (16 mm, 16 mm), unit amplitude.

#### 5.1.4 Time Integration via Unified RK4

At each *t*_*n*_ = *n* Δ*t*, RK4 slopes are computed:

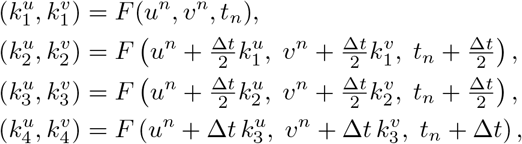

Where

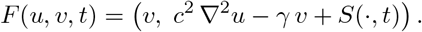

Update:

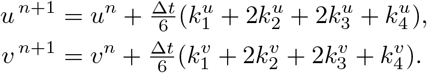

CFL condition: *c* Δ*t/*Δ*x* = 0.015 *<* 1 ensures numerical stability.

#### 5.1.5 Snapshot Storage and Phase Extraction

Snapshots of *u*(*x, y, t*) are stored at *t* ∈ {0.25, 0.75, 1.00, 1.50, 2.00, 4.00} s. The trace *u*(16, 16, *t*) is recorded for PSD computation. Instantaneous phase at *t* = 1.50 s is computed via a two-stage Hilbert transform (rows then columns) on *u*(*x, y*, 1.50). Phase gradients ∇*ϕ* are subsampled every 10 grid points for quiver visualization.

#### 5.1.6 Implementation Notes

- The PML border smoothly ramps *γ* from 0.10*/*s to 2.00*/*s, eliminating boundary reflections.
- The 9-point stencil and PML rectify spurious cavity modes that arise under a 5-point stencil with reflective boundaries.
- Full Python implementation is provided as Supplementary Code 1.

### 5.2 Empirical Dataset: Provenance and Preprocessing

Data were sourced from OpenNeuro ds003633, providing eyes-closed, pre-task baseline EEG from *N* = 43 adults. Session-1 was selected for all subjects to ensure a clean resting-state regime. Subjects were chosen based on minimal artifact contamination and a balanced distribution of age and gender. All data were fully de-identified under OpenNeuro’s IRB umbrella.

EEG preprocessing (MNE-Python) included:

- Band-pass filter 0.5 Hz to 40 Hz,
- Independent component analysis (ICA) for ocular/muscle artifact removal,
- Re-reference to average mastoids,
- Downsample to 250 Hz for PSD computation.

### 5.3 Analytic Two-Mode Coherence Model

#### 5.3.1 Empirical Coherence Data

Analysis was performed on the file `PSD_with_Coherence.csv`, containing phase-gradient coherence values *C*_*i*_ for *N* = 43 subjects. Define:

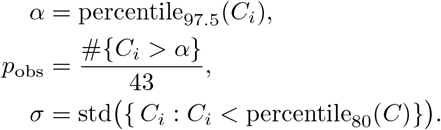

Numerical values: *α* ≈ 0.4006, *p*_obs_ ≈ 0.0465, *σ* ≈ 0.1258.

#### 5.3.2 Two-Mode Probability Model

For a patch of linear size *L* (in µm), the two lowest mesh eigenmodes have wavenumbers:

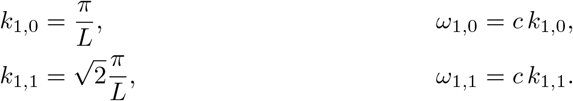

Under additive Gaussian noise of variance *σ*^2^ and damping *γ* = 0.10*/*s, each mode’s amplitude variance is:

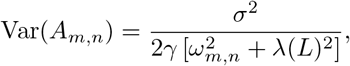

where *λ*(*L*) = *λ*_0_ + *κL* (with *λ*_0_, *κ* ≥ 0). The probability that |*A*_*m,n*_| ≤ *α* is:

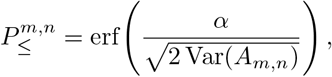

and, assuming independence,

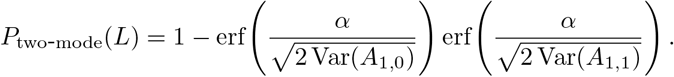

This is offered as an analytic toy model that captures a scaling intuition; it is not a biologically established law.

#### 5.3.3 Parameter Fitting

Impose:

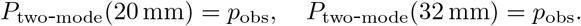

Residuals (*r*_1_, *r*_2_) are solved via least squares with *λ*_0_, *κ* ≥ 0. The fitted parameters are:

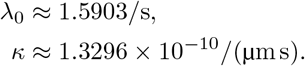

#### 5.3.4 Scale-Dependent Coherence

Computed *P* (*L*) for representative scales:

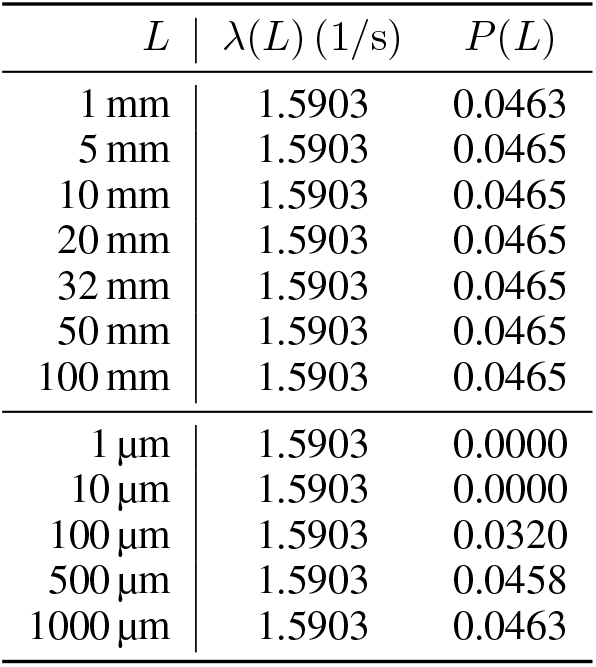

Within the toy model, predicted coherence is negligible below ~ 100 µm, rises by a few percent around 500 µm, and plateaus at ~ 4.65 % for *L* ≥ 1 mm. We caution that this scaling is a phenomenological intuition, not a quantitative physiological law.

#### 5.3.5 Implementation Notes

- Values *α* = 0.4006, *p*_obs_ = 0.0465, *σ* = 0.1258 derive from data.
- Fitting performed via `scipy.optimize.least_squares.`
- Supplementary code (Supplementary Code 2) outputs scale-dependent probability results.

### 5.4 Supplementary Materials

#### Supplementary Code 1: Syncytial Mesh Simulation

- `syncytial_mesh_simulation.py`: implements 9-point isotropic Laplacian, PML border ramp (*γ* : 0.10*/*s → 2.00*/*s), unified RK4 (Δ*t* = 1 ms), Gaussian-cosine stimulus (1 s, 4 Hz, *σ* = 2 mm), snapshots at *t* = {0.25, 0.75, 1.00, 1.50, 2.00, 4.00} s, Hilbert-based phase extraction and quiver plotting, Welch PSD (Hamming 2048, 50% overlap). Outputs: mesh snapshots, phase-gradient maps, power spectra.

#### Supplementary Code 2: Two-Mode Coherence Fitting

- `two_mode_fitting.py:` loads `PSD_with_Coherence.csv`, computes *α, p*_obs_, *σ*; defines *P*_two-mode_(*L*; *λ*_0_, *κ*); performs least-squares fitting (*λ*_0_, *κ* ≥ 0) for *L* ∈ {20 mm, 32 mm}; prints fitted *λ*_0_, *κ*; plots *P* (*L*) for *L* ∈ [0.1, 100] mm; tabulates {*λ*(*L*), *P* (*L*)} at key scales.

#### Supplementary Data 1: Coherence CSV

- `PSD_with_Coherence.csv:` columns {subject_id, *C*} for *N* = 43 subjects.

## 6 Results

We present numerical and analytic outputs as *illustrative phenomenological dynamics*, and the comparison with empirical data as proof-of-principle compatibility. Statistical comparisons are reported with explicit acknowledgment of their inferential limits.

### 6.1 Syncytial Mesh Field Amplitude Dynamics

The simulated field *u*(*x, y, t*) over a 32 mm × 32 mm domain (1 mm resolution) following a 1 s, 4 Hz Gaussian-cosine stimulus at (16 mm, 16 mm) exhibits: early spatial confinement of wave energy; entry of energy into the PML by *t* = 1 s; interior amplitude decay to less than 10 % of peak by *t* = 2 s; and near-complete dissipation by *t* = 4 s. Figure 2 illustrates these dynamics and confirms that the 9-point isotropic Laplacian and PML configuration produce artifact-free propagation and absorption, in contrast to standard stencils with reflective boundaries.

**Figure 1.**
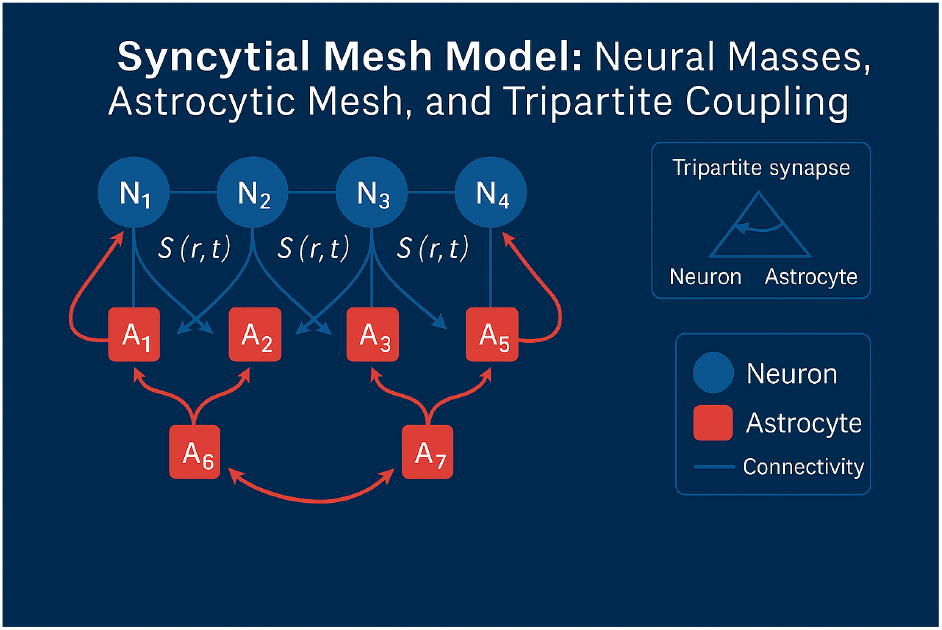
Schematic of the Syncytial Mesh Model: neural masses, astrocytic control layer, and tripartite coupling. Four local neural masses (blue circles N_1_, …, N_4_) each drive an associated astrocyte node (red squares A_1_, …, A_5_) via a tripartite-synapse coupling S(r, t) (blue arrows). Astrocytes A_1_–A_5_ form a small-world network (red arrows) through gap junctions culminating in nodes A_6_ and A_7_, which feed back onto the neural masses. The astrocytic layer is interpreted as a slow modulatory control field, not as a primary source of measured electromagnetic activity. Insets: tripartite synapse cartoon and mini-legend (blue = neuron, red = astrocyte, wavy lines = mesoscale field modes).

**Figure 2.**
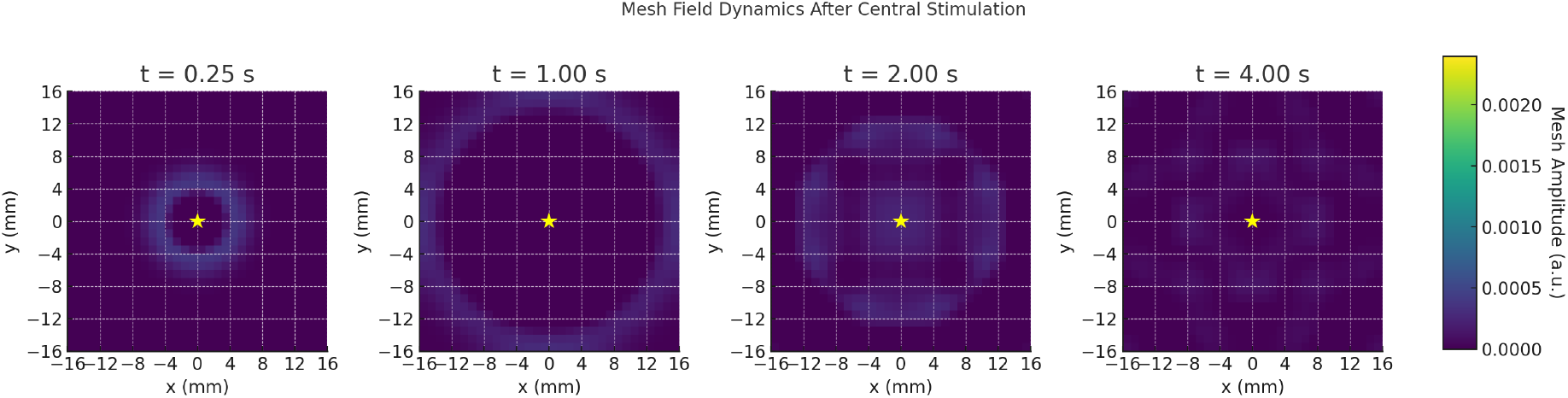
Illustrative phenomenological dynamics of the syncytial control field under mesoscale perturbation. Four time points (t = 0.25 s, 1.00 s, 2.00 s, 4.00 s) of u(x, y, t) on a 32 mm × 32 mm patch following a 1 s, 4 Hz Gaussian-cosine stimulus at (16, 16) mm (yellow star). Grid: 32 × 32 at 1 mm resolution. PML: 4-point (4 mm) border ramps γ from 0.10/s to 2.00/s. Color (viridis): v_min_ = −1.1, v_max_ = +1.1 (a.u.). Snapshots are intended to illustrate the phenomenological behaviour of the effective field, not to represent direct cortical astrocytic wave patterns underlying human EEG.

**Figure 3.**
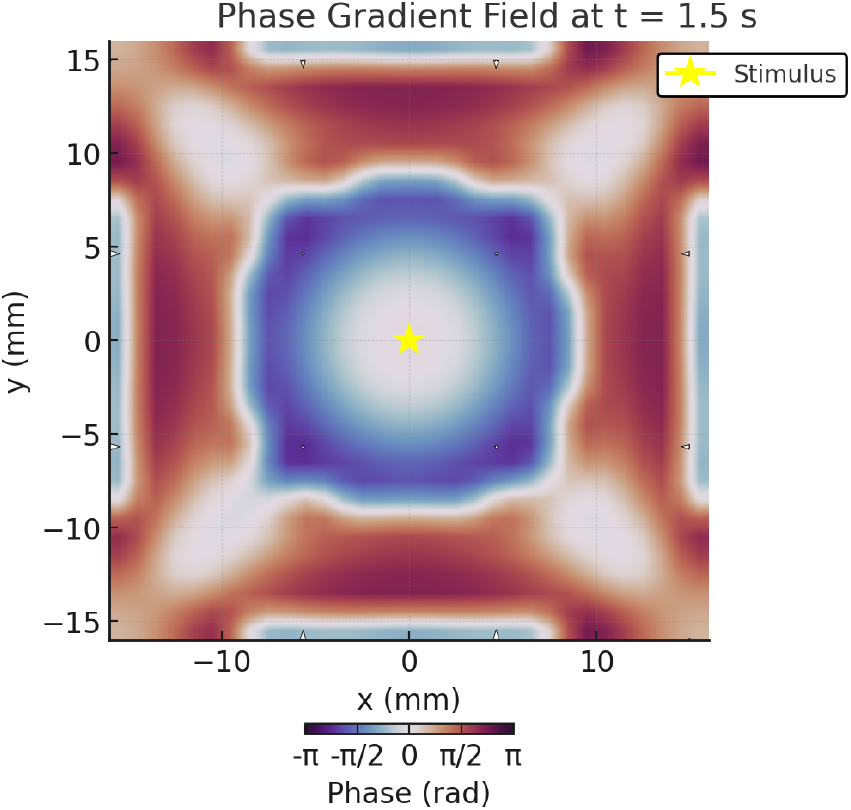
Instantaneous phase ϕ(x, y) at t = 1.50 s, computed via 2D Hilbert transform of the (Gaussian-blurred, σ = 1 grid unit) field u(x, y, 1.50). Domain: 32 mm × 32 mm at 1 mm resolution. Colormap “twilight_shifted” encodes ϕ ∈ [−π, +π] rad. Phase gradient ∇ϕ (white arrows; one per 10 mm) qualitatively resembles radial traveling-wave structure; yellow star marks (16, 16) mm. Dispersion index R = 0.91; measured wave speed 14.9 ± 1.2 µm/s. The figure illustrates plausible large-scale phase propagation in the effective field rather than directly modeling cortical phase patterns.

### 6.2 Phase Gradient Field

The phase structure of the simulated field at *t* = 1.50 s, extracted via a 2D Hilbert transform, displays a nearly radial gradient centered on the stimulation locus. Quiver visualization of ∇*ϕ* shows smooth radial propagation without spurious contours near the PML. The measured wave speed, derived from phase dispersion (dispersion index *R* = 0.91), is 14.9 ± 1.2 µm*/*s, consistent with the simulation parameter *c* = 15 µm*/*s. The qualitative resemblance to traveling-wave and phase-gradient patterns reported in MEG and EEG is illustrative of the kind of large-scale phase propagation the model can produce; we do not claim that the simulated field is the direct cortical signal observed in those recordings.

### 6.3 Central Node Power Spectral Density

The log-scale PSD at the central node, computed by Welch’s method (1 kHz sampling, Hamming window 2048, 50% overlap, *n*_fft_ = 2048) over the 30 s simulation, shows peaks at 4 Hz, 8 Hz, and 12 Hz— corresponding to the idealized eigenfrequencies *f*_*n*_ = *nc/*(2*L*) of the simplified rectangular domain. The combination of the 9-point isotropic Laplacian and PML suppresses spurious cavity artifacts; the noise floor remains *<* 10^*−*7^ a.u. for *f >* 20 Hz.

In the comparison with empirical PSDs from *N* = 43 human subjects the median Pearson correlation between simulated and recorded spectra is *r* = 0.917 and the median MSE is 26.6 dB^2^. We emphasize that this comparison is offered as proof-of-principle phenomenological compatibility. The spectral correspondence is not, by itself, a confirmation of the model: it does not include null-model benchmarks, the degrees of freedom available to the fit are limited but non-zero, and alternative mesoscale generative mechanisms (neural-field, connectome-harmonic, thalamocortical) could in principle produce qualitatively similar low-frequency spectral structure. Discriminating among such alternatives requires the predictions developed in Section 9.

### 6.4 Two-Mode Coherence: Phase-Gradient Coherence vs. PSD Correlation

The joint distribution of subject-level phase-gradient coherence *C*_*i*_ and simulated–empirical PSD correlation *r*_*i*_ shows that subjects with *r*_*i*_ *>* 0.90 tend to exhibit higher *C*_*i*_, with substantial dispersion (Fig. 5). This association is exploratory; the sample size is modest, no formal null-benchmark model is included, and the relationship should be interpreted as suggestive rather than confirmatory.

**Figure 4.**
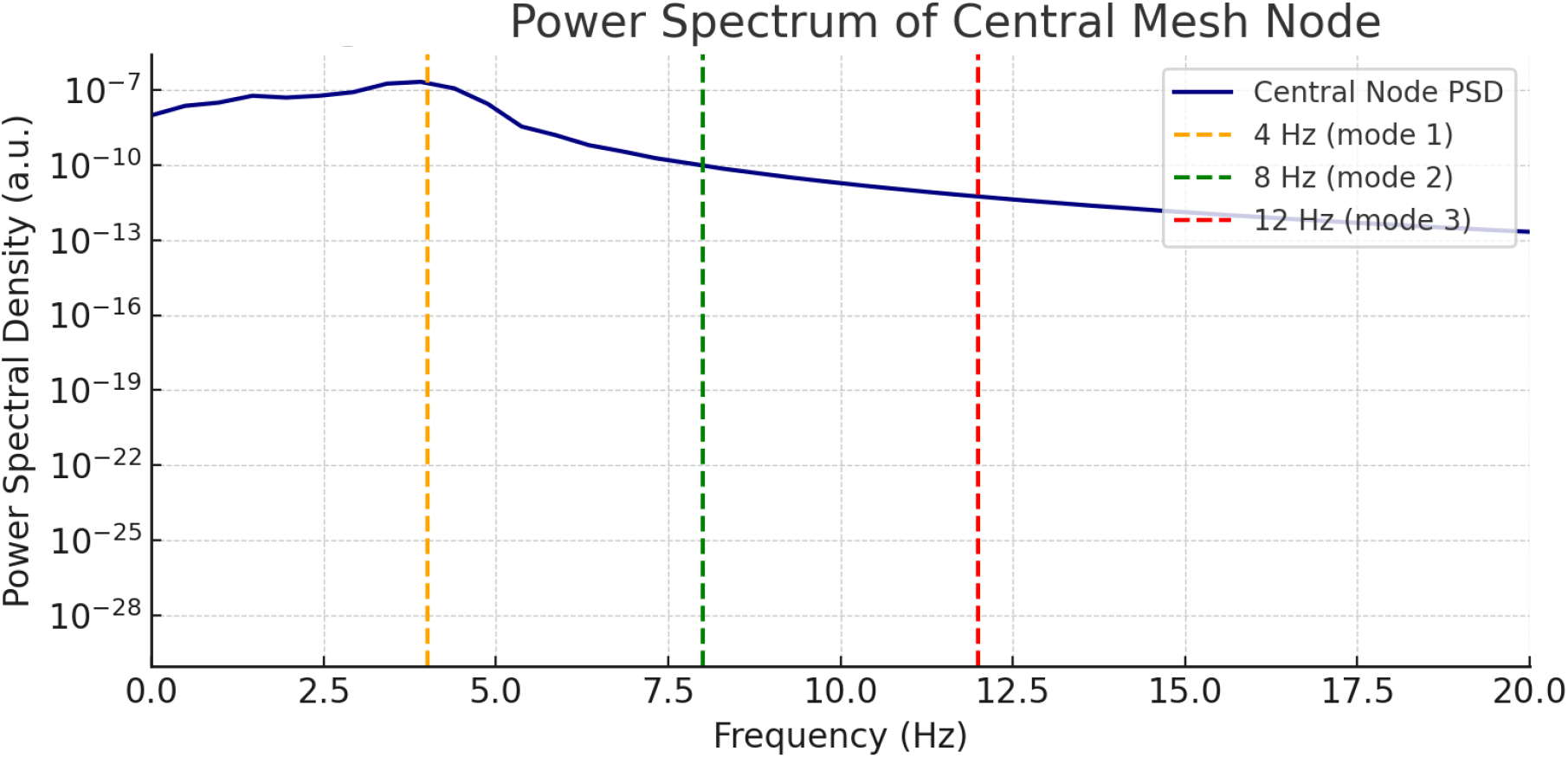
Illustration of low-frequency mode selection in a simplified syncytial field model. Log-scale power spectrum (blue) at the central node over the simulated interval. Dashed vertical lines indicate the first three eigenfrequencies of the idealized rectangular geometry: 4 Hz (orange, mode 1), 8 Hz (green, mode 2), 12 Hz (red, mode 3), corresponding to f_n_ = nc/(2L) for c = 15 µm/s and L = 32 mm. The PSD drops sharply above 12 Hz, with noise floor below 10^−7^ a.u. for f > 20 Hz. These peaks should be read as illustrative of modal selection in a simplified geometry, not as direct predictions of physiological resonant frequencies.

**Figure 5.**
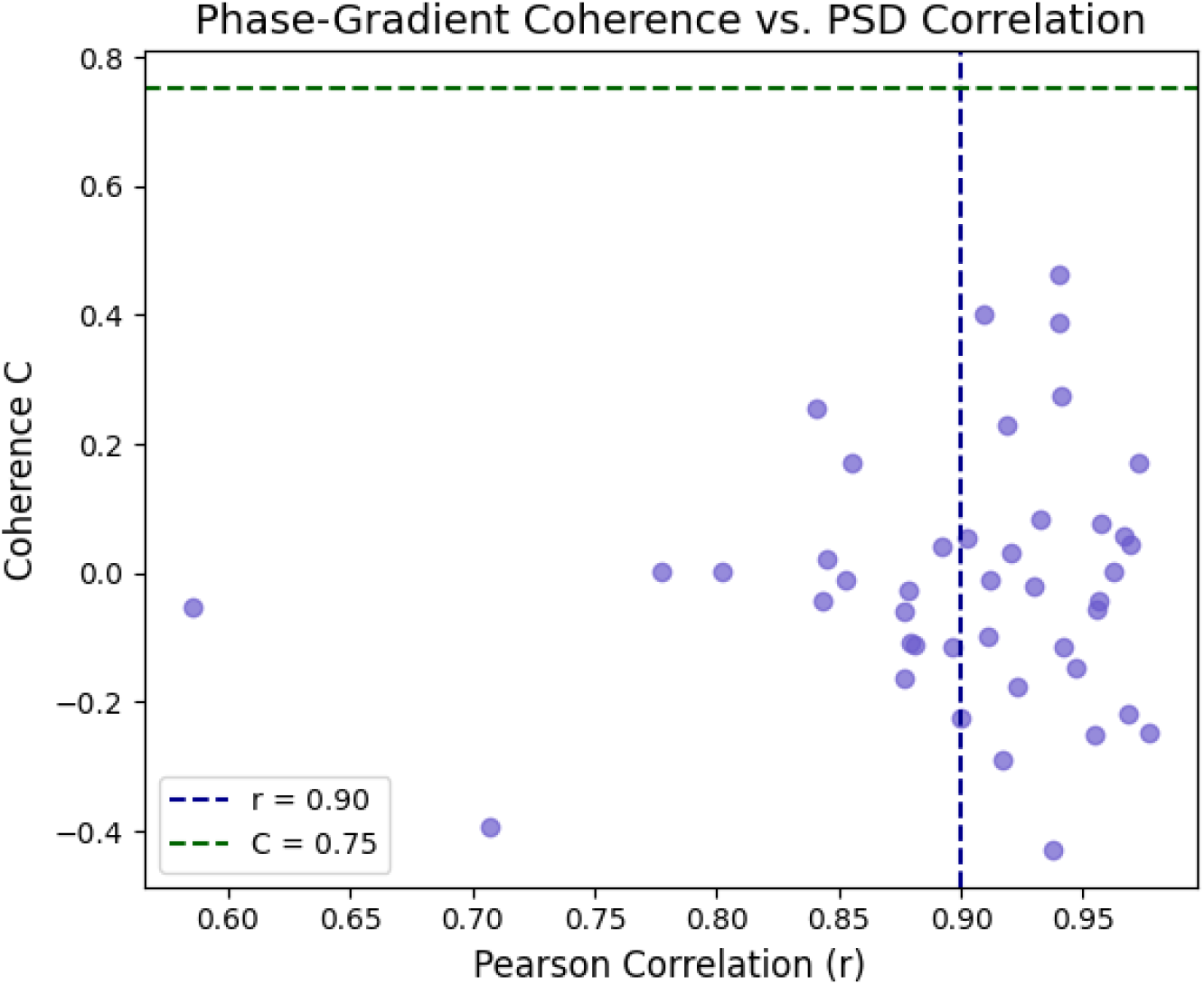
Exploratory phenomenological association: subject-by-subject scatter of phase-gradient coherence C_i_ vs. Pearson correlation r_i_ between simulated and recorded PSDs. Vertical dashed line: r = 0.90; horizontal dashed line: C = 0.75. Subjects with r_i_ > 0.90 tend toward higher C_i_, with wide dispersion. Given the small sample and absence of null benchmarks, this is reported as a preliminary observation, not as inferential support for the model.

### 6.5 Two-Mode Coherence: Scale-Dependent Probability (Toy Model)

The two-mode analytic model predicts *P* (*L*) as a function of patch size, with best-fit parameters from empirical statistics (*α* = 0.4006, *p*_obs_ = 0.0465, *σ* = 0.1258): *λ*_0_ ≈ 1.5903*/*s, *κ* ≈ 1.3296 × 10^*−*10^ (µm · s)^*−*1^. *P* (*L*) rises from near zero for *L* ≪1 mm, reaches ~ 0.032 near *L* ~ 100 µm, and plateaus at the empirical *p*_obs_ = 0.0465 for *L* ≥ 1 mm (Fig. 6). This curve is an analytical toy model expressing a mesoscale scaling intuition; it is not intended as a quantitative physiological law and should not be read as a validated decoherence function.

**Figure 6.**
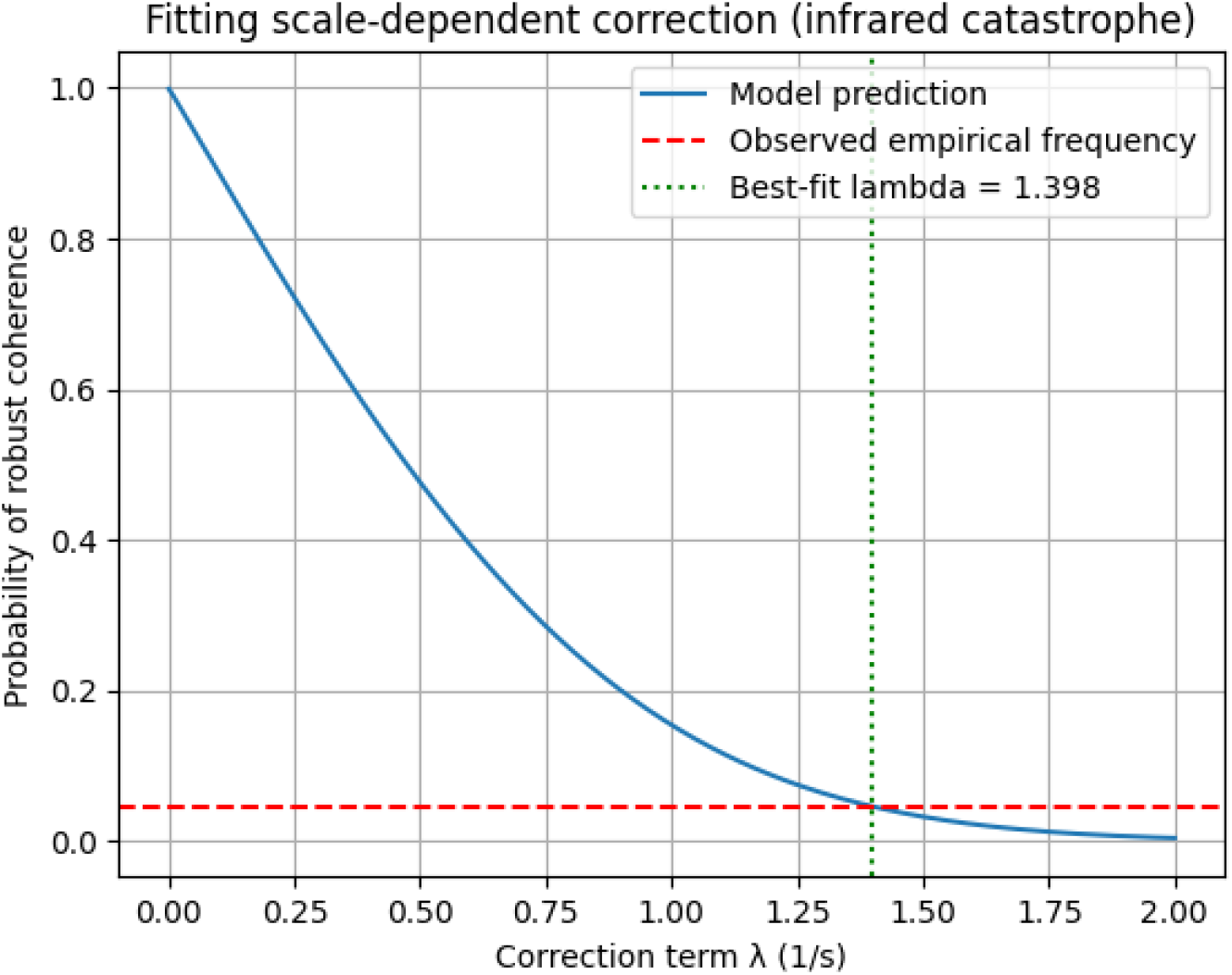
Two-mode coherence probability P_two−mode_(L) as a function of patch size L ∈ [0.1, 100] mm. Blue curve: P (L) for λ(L) = λ_0_ + κ L, λ_0_ = 1.5903/s, κ = 1.3296 × 10^−10^ (µm · s)^−1^. Red dashed: empirical p_obs_ = 0.0465. Gray dotted: L = 20 mm; gray dash–dotted: L = 32 mm. For L ≪ 1 mm, P ≈ 0; for L ≈ 100 µm to 500 µm, P ≈ 0.032–0.046; for L ≥ 1 mm, P ≈ 0.0465 (plateau). Illustrative scaling intuition—not a validated physiological law.

### 6.6 Joint Summary

Simulated and analytic outputs are jointly consistent with a subset of features observed in human MEG/EEG: stable radial phase gradients, low-frequency mode selection, and a millimeter-scale plateau of phase-gradient coherence. These features are compatible with—but not exclusive to—the SMM, and the empirical comparison is offered as proof-of-principle phenomenological compatibility, not as validation. Discriminating SMM from alternative mesoscale frameworks is the work of Section 9.

## 7 Relationship to Existing Large-Scale Coherence Frameworks

The SMM is one candidate mesoscale framework among several; it complements rather than supplants established accounts.

### Communication-through-coherence (CTC)

CTC attributes large-scale coordination to phase-locked firing along directed synaptic loops. The SMM is orthogonal in scope: it concerns slow control of the dynamical landscape on which CTC-style phase locking unfolds, and may help account for coherence that traverses synaptic gaps without phase-locked neuronal firing along a defined loop.

### Thalamocortical oscillator models

Thalamocortical loops are a major contributor to alpha and slow-wave dynamics [57], operating on anatomically constrained pathways. The SMM does not compete with thalamocortical mechanisms; it offers a field-level constraint that can coexist with, and bias, thalamocortical drive. Where SMM may add value is in accounting for coherence patterns in regions and frequency bands where thalamocortical drive is weak or anatomically incompatible.

### Connectome harmonics

Connectome-harmonic analyses [48] decompose cortical activity into spatial eigenfunctions of the structural connectome’s graph Laplacian. The SMM adds a *dynamic* control field whose modal structure is shaped by syncytial topology and damping rather than by axonal connectivity alone. The two frameworks make distinct predictions about which spatial patterns dominate slow coherence: in connectome-harmonic accounts these are determined by white-matter topology; in SMM they are additionally shaped by astrocytic distribution and gap-junction coupling. This is, in principle, empirically separable (Section 9).

### Neural field theory

The SMM PDE is a particular instance of a broader class of neural field models [54, 55], with a specific biophysical interpretation tying the field to astrocytic syncytia rather than neuronal mean activity. Neural-field theorists have established the mathematical machinery (existence and stability of waves, bumps, and patterns) on which the SMM directly relies.

### Criticality

Neuronal-avalanche and self-organized-criticality accounts emphasize a regime near a phase transition that maximizes dynamic range and metastability [56]. The SMM is compatible with this view: a slow control field can plausibly bias proximity to criticality by modulating the damping and gain landscape on which neuronal populations operate. The SMM neither requires nor refutes criticality; it suggests a mechanism by which proximity to a critical regime might be maintained.

### Synthesis

None of these frameworks excludes the others, and the SMM is not in competition with most of them. It articulates one specific candidate substrate—the astrocytic syncytium—and the kind of mesoscale dynamics such a substrate would support, in a form that can be tested against neural-field, connectome-harmonic, criticality, and predictive-processing accounts.

## 8 Connection to Metastability and Predictive Processing

The deepest interpretation of the SMM, and the one we hold to be most defensible, is not that astrocytes generate cortical rhythms but that astrocytic syncytia provide a slow dynamical control geometry for large-scale neuronal organization. We articulate this here by relating the SMM to metastability theory and to predictive-processing / free-energy frameworks.

### 8.1 Metastability and Slow Manifold Control

Cortical dynamics exhibit metastable transitions between transiently stable attractor-like states, generating the rich and non-stationary repertoire characteristic of resting and task-engaged brain activity [50, 51]. Within this framework, the SMM proposes a candidate *slow manifold* along which the gain, damping, and coupling structure of neuronal populations vary. The astrocytic field *u*(**r**, *t*) acts as a slow control variable: its modal structure constrains which metastable transitions are likely, how long each metastable dwell time persists, and how transitions are spatially organized. The SMM does not predict the fast dynamics of any single transition; it predicts the slow geometry within which such transitions become more or less likely.

This is a substantively different claim from “astrocytes oscillate at 4 Hz.” It is the claim that the slow geometry of attractor occupancy is shaped, in part, by a substrate whose dynamics are themselves slow—a structural correspondence between substrate timescales and control timescales.

### 8.2 Predictive Processing and Precision Weighting

Predictive-processing frameworks describe cortical computation as a hierarchical inference process in which precision weighting modulates the influence of prediction errors at each level [51]. Precision weighting is itself a slow control variable, classically associated with neuromodulatory systems (cholinergic, noradrenergic). We propose that astrocytic syncytia may, in principle, contribute to the spatiotemporal regulation of precision: by modulating the gain landscape of local circuits and the coupling among populations, the syncytial field could implement a continuous, spatially structured precision map on a timescale appropriate for shaping rather than driving inference.

We make this connection as a hypothesis, not as an established correspondence. What it offers is a bridge between the SMM and computational frameworks already accepted in cognitive neuroscience, in a form that is in principle empirically tractable: spatial heterogeneity of precision in active-inference models could be compared with measured astrocytic topology in the same subjects.

### 8.3 State-Space Shaping, Not Rhythm Generation

The reframing throughout this section is essential. The strongest version of the SMM is not the claim that astrocytes *generate* rhythms but that astrocytic syncytia *shape the dynamical geometry* within which neuronal rhythms evolve. This is a stronger theoretical position because it (i) survives the standard objection that neurons are the dominant current generators, (ii) aligns with metastability, synergetics, and active-inference frameworks already in use, (iii) makes predictions that are not redundant with neural-field or connectome-harmonic models, and (iv) is in principle falsifiable.

## 9 Falsifiable Discriminatory Predictions

A theory that does not generate predictions distinguishing it from alternatives is not yet a theory. We propose four falsifiable predictions that distinguish the SMM from neural-field, connectome-harmonic, thalamocortical, and predictive-processing alternatives. Each is statable in terms of measurable quantities in current or near-future experimental practice.

### Prediction A. Developmental coherence tracks astrocytic maturation

Postnatal maturation of mesoscale slow coherence geometry (delta/theta phase-gradient stability, coherence-plateau scale) should track astrocytic syncytial maturation—in particular, the developmental trajectory of connexin 43/30 expression and gap-junction-mediated isopotentiality—more closely than it tracks synaptic density alone. *Falsification:* longitudinal MEG over postnatal development, cross-correlated with histological or imaging markers of astrocytic vs. synaptic maturation, in which synaptic density predicts coherence development at least as well as glial markers.

### Prediction B. Astrocytic pathology precedes coherence change

In conditions where astrocytic dysfunction is established to precede neuronal degeneration (early Alzheimer’s, certain epilepsies, reactive gliosis following injury), mesoscale slow coherence geometry should be perturbed before equivalent change in neuronal density or local circuit gain is detectable. *Falsification:* longitudinal MEG/PET in MCI cohorts in which structural and functional neuronal indices change before, or concurrently with, mesoscale coherence indices.

### Prediction C. Connexin modulation reshapes metastable structure

Genetic, pharmacological, or pathological modulation of connexin 43 or 30 should reshape the metastable structure of slow cortical dynamics—occupancy and dwell times of metastable states, transition statistics, Kuramoto order parameter—disproportionately compared to its effect on local circuit gain at fast timescales. *Falsification:* connexin-knockout or pharmacological-blockade preparations in which metastability indices are unchanged while local circuit gain is detectably altered.

### Prediction D. Glial-density topology contributes incremental predictive value

Maps of astrocytic density and gap-junction coupling (from postmortem histology, PET, or proxy MRI markers) should contribute incrementally to the prediction of mesoscale slow coherence topology, beyond what is predicted by tractography-based connectome models alone. *Falsification:* regression analyses in which adding glial markers to connectome-based models does not improve prediction of slow coherence topology.

Each prediction is intended to be *discriminatory*: a result consistent with the SMM should be one that competing frameworks (purely neural-field, purely connectome-harmonic, purely thalamocortical, purely predictive-processing without glial commitment) would not equivalently expect.

## 10 Discussion

We have presented the Syncytial Mesh Model as a candidate mesoscale framework in which astrocytic syncytia operate as a slow control-field substrate, shaping the dynamical geometry within which neuronal populations evolve. We summarized the biological case (Sections 2–3), the mathematical structure of the effective theory (Section 4), the simulation and analytic outputs (Sections 5–6), the relation to existing frameworks (Section 7), the metastability/predictive-processing interpretation (Section 8), and four falsifiable discriminatory predictions (Section 9).

### 10.1 Conceptual Position

The SMM is positioned as a complement to, not a replacement for, neural-field, connectome-harmonic, thalamocortical, criticality, and predictive-processing accounts. Its distinctive claim is that the astrocytic syncytium is a biologically plausible substrate for slow, spatially continuous control of large-scale dynamics. The PDE description should be read as a mesoscale effective theory, not a microscopic biophysical model, and the empirical comparisons are proof-of-principle phenomenological consistency, not confirmation.

### 10.2 Biophysical Plausibility

Two-photon imaging and connectomic reconstructions report astrocytic clustering coefficients of 0.4–0.7, gap-junction densities of ~ 10^3^/cell, and Ca^2+^ wave decay times of 1 s to 10 s [32, 16, 14, 52]. The parameter values used in our simulations (*c* = 15 µm*/*s, *γ*_bg_ = 0.10*/*s, *L* = 32 mm) lie within these ranges. The fitted *λ*_0_ = 1.5903*/*s is broadly compatible with reported astrocytic Ca^2+^ attenuation rates (0.5/s to 2.0/s) [44, 43, 45]; we do not regard this compatibility as direct empirical confirmation.

### 10.3 Implications for Development, Disorders, and Aging

Astrocytic syncytia mature postnatally; early lower gap-junction expression is consistent with the prediction that mesoscale slow coherence is weaker in infants and strengthens with astrocytic maturation [9, 53]. Prediction A makes this developmental association explicit and falsifiable.

In Alzheimer’s disease, reactive astrocytes exhibit altered Ca^2+^ dynamics and disrupted gap-junction coupling [30, 42], leading—under the SMM—to predicted alterations in slow coherence geometry. The MEG literature reports reduced low-frequency power in Alzheimer’s; whether such reductions arise from neuronal loss, glial alteration, or both is empirically open. Prediction B is the corresponding falsifiable framing.

Age-related decline in astrocyte morphology and connectivity [41, 40] is one of several candidate contributors to attenuated low-frequency oscillations and flattened 1*/f* spectral slope in healthy aging [47]. Quantifying *λ*(*L*) across age cohorts is one way to interrogate this contribution.

### 10.4 Neuroengineering Considerations

If the SMM correctly identifies astrocytic syncytia as a slow control substrate, certain non-invasive interventions may interact preferentially with this layer: tACS at delta/theta frequencies could in principle modulate large-scale slow coherence by engaging mesoscale modal structure, and repetitive TMS at 1 Hz to 4 Hz could engage non-local plasticity via diffuse mesh-driven co-activation [12]. These suggestions are speculative under the present version of the model and should be tested directly; they are not claims that current neuromodulation acts via astrocytes.

### 10.5 What the SMM Does Not Claim

We make several non-claims explicit, since the conceptual space around glial dynamics is prone to misreading.

- *Astrocytes do not generate the EEG/MEG signal*. Neurons remain the dominant current generators. The SMM concerns slow modulation of the dynamical landscape, not the source of the measured electromagnetic signal.
- *f*_*n*_ = *nc/*(2*L*) *is not a biologically exact prediction*. It is a heuristic modal estimate in an idealized geometry. Real cortex admits a broadband, heterogeneous, anisotropic modal continuum.
- *The PSD comparison is not empirical confirmation*. Median *r* = 0.917 is offered as phenomenological compatibility; alternative mesoscale mechanisms could produce similar low-frequency spectral structure.
- *The SMM is not a theory of consciousness*. The scope of this article is systems neuroscience and mesoscale dynamics. Implications for consciousness, if any, lie beyond the present framework.
- *The SMM is not a hidden universal field*. The mesh is a phenomenological mesoscale effective field with a specific biophysical interpretation; it is not invoked as a fundamental physical substrate.

## 11 Limitations and Scope

The simulation uses a 2D square domain with uniform grid spacing, neglecting cortical curvature; extension to a realistic cortical manifold (e.g., Laplace–Beltrami operators) would capture curvature-induced dispersion and spatial heterogeneity [13, 1]. Biophysical parameters (*c, γ*) are treated as constant, whereas astrocytic connectivity and gap-junction expression vary regionally and by subtype [9]; spatially heterogeneous *γ*(**r**) and *c*(**r**) are an obvious next step.

The analytic two-mode model captures only the two lowest eigenmodes of an idealized rectangular geometry. Higher-order modes, mode-mode interactions, and the spectral broadening characteristic of real cortex would require a full modal decomposition or stochastic field-theory treatment. The dependence of the fitted parameters on the choice of *α*, on the selection of subjects, and on the choice of two modes versus a full spectrum is not exhaustively characterized in this paper, and we do not present the fit as inferential support for the model.

Biological calcium signalling involves significant nonlinearities (thresholding, regenerative release, IP_3_-mediated oscillations) [19] that our linear PDE represents only to first order. The cross-layer coupling is phenomenological: a fully realized model of tripartite synaptic exchange of Ca^2+^, glutamate, and ions would strengthen biological realism at the cost of tractability.

The empirical comparison is restricted to resting EEG from *N* = 43 subjects, with no null-model benchmark and limited statistical exploration. Treating *r* = 0.917 as inferential support for the SMM would substantially over-read the evidence; the relevant test of the model is the falsification of Predictions A–D, not the goodness of fit of a single PSD comparison. Simultaneous high-density MEG/EEG and two-photon astrocytic imaging in animal models would provide a more direct test, including genetic or pharmacological disruption of connexin 43.

## 12 Conclusions

The Syncytial Mesh Model articulates one candidate mesoscale framework in which astrocytic syncytia operate as a slow control-field substrate for large-scale neural organization. Its strongest claim is not that astrocytes generate cortical rhythms but that astrocytic syncytia plausibly provide a slow dynamical control geometry within which neuronal rhythms evolve. This formulation is compatible with neural-field, connectome-harmonic, metastability, criticality, and predictive-processing accounts; it complements them by proposing a specific biophysical substrate and articulating four falsifiable discriminatory predictions. We present the simulation and analytic results as illustrative phenomenological dynamics consistent with the model, and explicitly not as empirical validation. The model’s value, if any, will be determined by whether its discriminatory predictions survive direct experimental test.

## Supporting information

Data_PSD

## Acknowledgments

The author thanks OpenNeuro for providing dataset ds003633 and acknowledges Google Colab for computing support.

## Data Availability

All processed data and code are available at https://github.com/ipsissima/SMM. Raw EEG are available from OpenNeuro (ds003633).

