## Supplementary material for "The Syncytial Mesh Model: A Mesoscale Control-Field Framework for Scale-Dependent Coherence in the Brain": Data_PSD: compare_psd.py

```
"""
Compare simulated PSD vs all EEG PSD CSVs. Outputs summary.
Contact: Andreu.Ballus@uab.cat
"""
import os
import pandas as pd
import numpy as np
from scipy.stats import pearsonr
from sklearn.metrics import mean_squared_error

sim_psd_path = "Figure4_PSD.csv"
eeg_psd_folder = "/content/drive/MyDrive/Ongoing projects/PSD_Results"

sim_df = pd.read_csv(sim_psd_path)
sim_freqs = sim_df['frequency_Hz'].values
sim_psd = 10 * np.log10(sim_df.iloc[:, 1].values + 1e-12)

results = []
for fname in os.listdir(eeg_psd_folder):
    if not fname.endswith('.csv'): continue
    df = pd.read_csv(os.path.join(eeg_psd_folder, fname))
    eeg_freqs = df['frequency_Hz'].values
    eeg_psd = df[df.columns[1]].values
    if 'dB' not in df.columns[1]:
        eeg_psd = 10 * np.log10(eeg_psd + 1e-12)
    eeg_psd_interp = np.interp(sim_freqs, eeg_freqs, eeg_psd)
    corr, _ = pearsonr(sim_psd, eeg_psd_interp)
    mse = mean_squared_error(sim_psd, eeg_psd_interp)
    results.append({'subject': fname, 'corr': corr, 'mse': mse})
results_df = pd.DataFrame(results)
results_df.to_csv('PSD_Comparison_Summary.csv', index=False)
print(results_df.describe())
```
