## Supplementary material for "The Syncytial Mesh Model: A Mesoscale Control-Field Framework for Scale-Dependent Coherence in the Brain": Data_PSD: plot_psd_comparison_results.py

```
"""
Plot summary statistics of PSD comparisons.
Contact: Andreu.Ballus@uab.cat
"""
import pandas as pd
import matplotlib.pyplot as plt
import seaborn as sns

df = pd.read_csv('PSD_Comparison_Summary.csv')
plt.figure(figsize=(8, 4))
sns.histplot(df['corr'], bins=20, kde=True)
plt.title('Distribution of PSD Correlation (simulated vs EEG)')
plt.xlabel('Pearson r')
plt.tight_layout()
plt.savefig('corr_hist.png', dpi=150)
plt.close()
plt.figure(figsize=(8, 4))
sns.histplot(df['mse'], bins=20, kde=True)
plt.title('Distribution of MSE (simulated vs EEG)')
plt.xlabel('Mean Squared Error')
plt.tight_layout()
plt.savefig('mse_hist.png', dpi=150)
plt.close()
print(df.describe())
```
