## Supplementary material for "The Syncytial Mesh Model: A Mesoscale Control-Field Framework for Scale-Dependent Coherence in the Brain": Data_PSD: preprocess_eeg_psd.py

```
"""
Preprocess EEG BIDS dataset, compute PSDs, save one CSV per subject.
Requires: mne, mne-bids, numpy, pandas, matplotlib.
Contact: Andreu.Ballus@uab.cat
"""
import os
import numpy as np
import pandas as pd
from mne_bids import BIDSPath, read_raw_bids

bids_root = '/content/drive/MyDrive/Ongoing projects/Neurodata'
output_dir = '/content/drive/MyDrive/Ongoing projects/PSD_Results'
os.makedirs(output_dir, exist_ok=True)

subject_list = [f"{i:03d}" for i in range(1, 44)]  # Subjects '001' to '043'
for subj in subject_list:
    try:
        bids_path = BIDSPath(subject=subj, session='1', task='EyesOpen',
                             acquisition='pre', datatype='eeg', root=bids_root)
        raw = read_raw_bids(bids_path=bids_path, verbose=False)
        raw.load_data().pick_types(eeg=True)
        raw.filter(1., 40., fir_design='firwin')
        raw.set_eeg_reference('average', projection=True)
        raw.resample(500)
        raw.crop(tmax=30)
        psd = raw.compute_psd(fmin=0.5, fmax=20, n_fft=2048, n_overlap=1024, n_per_seg=2048)
        freqs = psd.freqs
        psds_db = 10 * np.log10(psd.get_data() + 1e-12)
        mean_psd = psds_db.mean(axis=0)
        df_psd = pd.DataFrame({'frequency_Hz': freqs, 'PSD_dB': mean_psd})
        df_psd.to_csv(os.path.join(output_dir, f"sub-{subj}_PSD.csv"), index=False)
        print(f"Done: sub-{subj}")
    except Exception as e:
        print(f"Error for sub-{subj}: {e}")
```
