## Supplementary material for "The Syncytial Mesh Model: A Mesoscale Control-Field Framework for Scale-Dependent Coherence in the Brain": Data_PSD: README.md

# SMM PSD Pipeline

Reproducible workflow for comparing Syncytial Mesh Model (SMM) simulation spectra with empirical EEG data.

**Contact:**

## Steps

1. Preprocess EEG to PSD: `python preprocess_eeg_psd.py`
2. Simulate mesh PSD: `python simulate_mesh_psd.py`
3. Compare: `python compare_psd.py`
4. Plot/visualize: `python plot_psd_comparison_results.py`

- All scripts are set for the Google Colab/MNE workflow and expect your drive structure as used above.
- Example CSVs and `PSD_with_Coherence.csv` can be added as supplementary data.

## Requirements

```
pip install mne mne-bids numpy pandas matplotlib seaborn scipy
```

---

## Citation
If you use this code, please cite the associated preprint.
