## Supplementary material for "The Syncytial Mesh Model: A Mesoscale Control-Field Framework for Scale-Dependent Coherence in the Brain": Data_PSD: simulate_mesh_psd.py

```
"""
Simulate Syncytial Mesh, output central node PSD as CSV.
Contact: Andreu.Ballus@uab.cat
"""
import numpy as np
import matplotlib.pyplot as plt
import pandas as pd

L = 32
dx = 1.0
nx = int(L/dx)
ny = nx
x = np.linspace(0, L, nx)
y = np.linspace(0, L, ny)
X, Y = np.meshgrid(x, y)

dt = 0.001
T = 30.0
nt = int(T/dt)
c = 0.015
gamma_bg = 0.1
gamma_pml = 2.0
pml_width = 4

gamma = gamma_bg * np.ones((ny, nx))
for i in range(nx):
    for j in range(ny):
        if i < pml_width or i >= nx - pml_width or j < pml_width or j >= ny - pml_width:
            gamma[j, i] = gamma_pml

u = np.zeros((ny, nx))
v = np.zeros((ny, nx))

def laplacian_9pt(u, dx):
    lap = (
        -20 * u
        + 4 * (np.roll(u, 1, axis=0) + np.roll(u, -1, axis=0)
             + np.roll(u, 1, axis=1) + np.roll(u, -1, axis=1))
        + (np.roll(np.roll(u, 1, axis=0), 1, axis=1) + np.roll(np.roll(u, 1, axis=0), -1, axis=1)
         + np.roll(np.roll(u, -1, axis=0), 1, axis=1) + np.roll(np.roll(u, -1, axis=0), -1, axis=1))
    ) / (6 * dx**2)
    return lap

def stimulus(t):
    stim_dur = 1.0
    stim_freq = 4.0
    stim_amp = 1.0
    sigma = 2.0
    if t > stim_dur:
        return np.zeros((ny, nx))
    xc, yc = L/2, L/2
    spatial = np.exp(-((X-xc)**2 + (Y-yc)**2) / (2*sigma**2))
    temporal = stim_amp * np.cos(2 * np.pi * stim_freq * t)
    return spatial * temporal

central_trace = []
for t_idx in range(nt):
    t = t_idx * dt
    S = stimulus(t)
    def rhs(u, v, t):
        lap = laplacian_9pt(u, dx)
        return (v, c**2 * lap - gamma * v + S)
    k1u, k1v = rhs(u, v, t)
    k2u, k2v = rhs(u + 0.5*dt*k1u, v + 0.5*dt*k1v, t + 0.5*dt)
    k3u, k3v = rhs(u + 0.5*dt*k2u, v + 0.5*dt*k2v, t + 0.5*dt)
    k4u, k4v = rhs(u + dt*k3u, v + dt*k3v, t + dt)
    u += (dt/6)*(k1u + 2*k2u + 2*k3u + k4u)
    v += (dt/6)*(k1v + 2*k2v + 2*k3v + k4v)
    central_trace.append(u[ny//2, nx//2])
central_trace = np.array(central_trace)
fs = int(1/dt)
from scipy.signal import welch
f, pxx = welch(central_trace, fs=fs, window='hamming', nperseg=2048, noverlap=1024, nfft=2048)
df_psd = pd.DataFrame({'frequency_Hz': f, 'PSD_a.u.': pxx})
df_psd.to_csv('Figure4_PSD.csv', index=False)
```
